# The role of optogenetic stimulations of parvalbumin-positive interneurons in the prefrontal cortex and the ventral hippocampus on an acute MK801 model of schizophrenia-like cognitive inflexibility

**DOI:** 10.1101/2022.03.25.485752

**Authors:** Enrico Patrono, Karolina Hrůzova, Jan Svoboda, Aleš Stuchlík

**Author notes:** ***Corresponding author*** Correspondence: Enrico Patrono, Lab. of Neurophysiology of Memory, Institute of Physiology CAS, Videnska 1083, 14200, Prague, Czech Republic. ***Contributions*** EP and KH conceived the study and performed experiments. EP performed data analysis and wrote the manuscript. AS and JS revised the manuscript. All authors read and approved the final manuscript.

## Abstract

**Background and Hypothesis:** Schizophrenia research arose in the last decades, focusing more on its neural basis. Executive functions such as decision making and cognitive flexibility are the main cognitive areas that are impaired and are considered schizophrenia endophenotypes. Recently, cognitive impairment has been connected with the ablation of glutamatergic NMDARs resulting in increased cortical activity. Selective NMDARs antagonists such as dizocilpine have been used to model cognitive inflexibility in schizophrenia. Moreover, a decreased GABAergic inhibitory activity has been shown elsewhere along with the enhanced cortical activity. This NMDARs/GABA unbalanced ratio may reduce the entrainment of prefrontal gamma and hippocampal theta rhythm, resulting in a prefrontal-hippocampal gamma/theta band desynchronization.

**Study Design:** The study addressed the role of acute administrations of dizocilpine to model schizophrenia-like cognitive inflexibility in rats. We used a new version of the attentional set-shifting task, where rats learned switching/reversing the relevant rule. Moreover, we used the new ASST after dizocilpine systemic injections to test cognitive flexibility. Finally, we used in vivo optogenetic stimulations at specific light pulses of parvalbumin-positive interneurons in the prefrontal cortex and ventral hippocampus.

**Results:** The first experiments showed that acute dizocilpine in rats reproduced schizophrenia-like cognitive inflexibility. The second set of experiments demonstrated that appropriate optogenetic light pulses frequencies could rescue the cognitive flexibility previously altered by acute dizocilpine.

**Conclusions:** These findings advance our knowledge on the pivotal role of parvalbumin interneurons in schizophrenia-like cognitive impairment and may serve as a standpoint for further research of this severe psychiatric disorder.

## Introduction

Schizophrenia (SCZ) is a severe psychiatric disorder affecting about 1% of the population, characterized by positive and negative symptoms (1). Executive functions are dramatically impaired in SCZ patients compared to healthy subjects (2) and are essential hallmarks and endophenotypes of SCZ (3-4). Cognitive flexibility refers to the ability to reverse/switch to a new relevant rule to solve a task (5). The attentional set-shifting task (ASST) is a behavioral task that has been validated to model cognitive flexibility in rats (6). It includes tasks where animals reverse/switch previous associations between sets of different stimuli.

Centrally, a functional relationship has been found between N-methyl-D-aspartate subtypes glutamatergic receptors (NMDARs) and gamma-aminobutyric acid (GABA), helping to explain the neurophysiological causes inducing SCZ-like cognitive impairments in the prefrontal cortex (PFC) and the hippocampus (HPC) (7-9). Moreover, studies have shown that NMDAR antagonists blocking the GluN2A subunit increase cortical excitatory activity in both PFC (10-12) and ventral HPC (vHPC), involving GABAergic interneurons (13). Furthermore, it has mainly been recognized that both prelimbic and infralimbic regions of PFC are connected with vHPC (14-15). Therefore the PFC subregions are well placed to monitor changing environmental contexts to engage in the orchestration of the most contextually-appropriate behavior through their efferent projection sites (16). Finally, inhibitory dysfunctions mediated by NMDARs may lead to maladaptive interactions between brain areas associated with crucial features of SCZ (17-18). Thus, NMDAR-binding drugs have been used to model the SCZ-like cognitive impairment in rodents, such as dizocilpine (MK801) (19). Furthermore, it is well-known that MK801 is more potent at NMDARs than phencyclidine (PCP), having different effects on other neurotransmitter systems, such as the dopaminergic and the serotoninergic ones (20). However, whether MK801 was used chronically, sub-chronically, or acutely, this affected the way to model SCZ-like cognitive impairments, especially cognitive flexibility (21-23). Nevertheless, it was demonstrated that acute MK801 treatment involved only the cognitive abilities investigated (reversal memory and cognitive flexibility) (24-25). Therefore, in this study, we sought to create a new model of SCZ-like cognitive inflexibility in young adult male rats using acute MK801 systemic injections during the reverse/switching rule sessions of a new version of the ASST.

Recently, it has been shown that both chronic (26) and acute (27) MK801 decreases the density of GABAergic parvalbumin-positive (PV+) interneurons, suggesting a mechanism by which an excitatory/inhibitory imbalance might occur in SCZ. Furthermore, in the last decade, several investigations described the critical role of NMDAR onto PV+ GABAergic interneurons for gamma oscillations and synchronization in cognitive functions (28-31), raising discussions on the possibility to enhance the GABA levels after MK801-induced NMDAR/GABA unbalanced ratio. That is because PV+ GABAergic interneurons can regulate firing rate, spike timing, and synchronization, allowing for information encoding in both the PFC and vHPC (32-33). Furthermore, it has been suggested that enhancement of theta/gamma coupling reflects a compensatory mechanism to maintain cognitive performance in the setting of increased difficulty (34-35). However, to our knowledge, there are no studies connecting directly flexible behavior in rats with a specific role of PV+ enhancement in PFC and vHPC and gamma/theta waves oscillation in a model of inflexibility induced by acute MK801 systemic injections.

Consequently, the second aim of this study was to evaluate the role of frequency-specific optogenetic stimulations of PV+ in PFC and vHPC in the acute MK801 model of SCZ-like cognitive inflexibility. We used light pulsing frequencies that resembled the gamma and the theta oscillations in PFC and vHPC, respectively, during the reverse/switching rule sessions of the ASST. The stimulation of PV+ at gamma or theta-like frequencies while performing a cognitive task is achievable only using in vivo optogenetics.

## Materials and Methods

All experiments and animal treatments complied with the Animal Protection Code of the Czech Republic and the European Community Council directive (2010/63/EC). Pilot data showed that our female rats presented high variability in performing the task independently from the estrous cycle’s period. Therefore, male Long-Evans rats (15-20 weeks old) were used for the first experiment. For the optogenetic experiments, the strain LE-Tg (Pvalb-iCre)2Ottc (PV-Cre) were created by Brandon Harvey and Jim Pickel (Optogenetics and Transgenic Technology Core, NIDA, USA), expressing Cre recombinase under the rat PV+ promoter, and kept at the Rat Resource and Research Center (RRRC, Columbia, MO, USA) (RRRC# 773). All rats were bred and housed in pairs with food and water available ad libitum and maintained on a 12/12 hr light-dark cycle. Animals were food-restricted and held at 85–90% of their body weight throughout the experiments. The experiments were performed during the light phase of the day.

### Drugs

(+)MK-801 (dizocilpine hydrogen maleate; SigmaAldrich, CR) was dissolved in 0.9% NaCl, at concentrations of 0.08 mg/ml, and injected intraperitoneally at 0.08 mg/kg. The specific dose was chosen to induce cognitive disabilities but not alter locomotion based on previous pilot data. Fresh solutions were prepared on the injection day.

The Cre-dependent viral vector pAAV-Ef1a-DIO -hChR2(E123T/T159C)-EYFP - serotype AAV9 - (ChR2) was a gift from Karl Deisseroth (Addgene plasmid # 35509; http://n2t.net/addgene:35509; RRID: Addgene_35509) (36).

### The attentional set-shifting task

The ASST is based on switching/reversing a discrimination rule to solve the task to be rewarded. Incorrect trials total number before reaching a criterion is counted for which the task is considered to be solved. In normal conditions, a rat can keep a low number of incorrect trials before reaching the criterion and therefore move to the following sessions. However, the rat alters the behavioral performance under drug treatment by increasing the number of incorrect trials to reach the criterion of reduced cognitive flexibility. Therefore, in the first set of experiments, we measured the ability to switch/reverse the relevant rule to solve the task using a new version of the ASST (37) (see supplementary materials).

Two groups of rats, experimental (MK801) (N=8) and control (NaCl) (N=8), were used. At the reversal and set-shifting sessions (CDrev, IDSrev, and EDS), the rats were acutely injected with MK801 or NaCl, respectively. MK801 was challenged (0.08mg/Kg, i.p.) according to preliminary results showing cognitive deficits but not locomotory alterations.

### The optogenetic stimulation of parvalbumin interneurons in the prefrontal cortex and ventral hippocampus

The second part of the study focused on stimulating PV+ interneurons in the PFC or vHPC during ASST and used the acute MK801 model to induce cognitive inflexibility using PV-Cre rats. Again, transgenic rats (heterozygous, HET) and wild-types (controls, WT) from the same litters were used for the experiments.

### Surgeries and viral injections

All rats weighed 300–350 g at the time of the surgery. Unilateral viral injections and optic fiber implants were pseudo-randomly balanced between the right and left hemispheres (see supplementary materials).

### Behavioral assessment

Three-four weeks after surgery, the groups of animals (PFC: HET [N=7], WT [N=7]; vHPC: HET [N=8], WT [N=8]) underwent the same behavioral protocol as in the previous set of experiments, except that the acute MK801 systemic injections were paired with optogenetic stimulation during the reversal, and the set-shifting sessions (see supplementary materials). In addition, we delivered a light pulsing at a gamma-like frequency (20ms, 50Hz, 5-7mW, 1min) for PFC experiments and a light pulsing at a theta-like frequency (50ms, 10Hz, 5-7mW, 1min) for vHPC experiments.

### Optic fiber position and viral expression evaluation

Immunohistochemical assays were performed to label PV+ and ChR2 co-localization (supplementary materials).

### Data analysis

Data were analyzed through ANOVA, and analyses involving repeated measures adopted a multivariate approach. Data in figures are represented as mean ± SEM unless otherwise stated. Statistical analysis was performed using GraphPad Prism 9.1.2. The level of significance was at α < 0.05. For further details, see supplementary materials.

## Results

*The ASST:* the ability to reverse/switch the previously positive pot association was evaluated in all groups (NaCl vs. MK801; PFC_HET vs. PFC_WT; vHPC_HET vs. vHPC_WT). A two-way RM ANOVA revealed an effect of the groups (F [5, 40]=22.65; p<0.0001), of the sessions (F [1.736, 69.44]=4.332; p=0.0212), and of the interaction (sessions x groups) (F [10, 80]=2.225; p=0.0242). Tukey’s multiple comparisons between the sessions revealed a significantly higher number of trials before reaching the 6-CCT in MK801, PFC_WT, and vHPC_WT groups with respect to the NaCl, PFC_HET, and vHPC_HET groups, particularly in the CDrev and EDS sessions (Fig 3a), suggesting that the heterozygous groups of both brain regions, if appropriately stimulated, resemble the NaCl group. Therefore, one can argue that the optogenetic stimulations of PV+ interneurons in the PFC and vHPC counteracted the previous MK801 acute injection, avoiding cognitive inflexibility.

**Fig 1:**
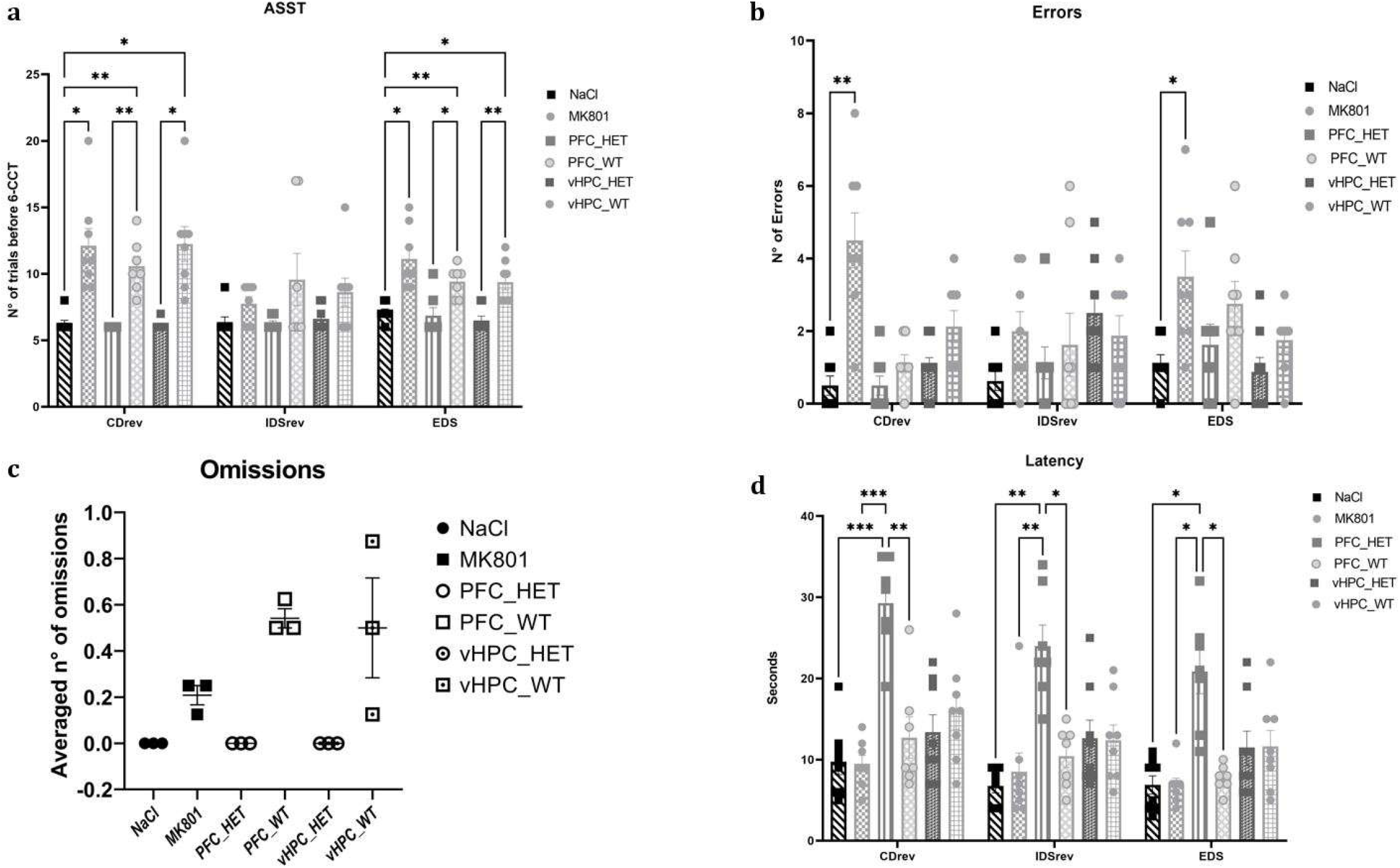
The ASST apparatus, optogenetic settings, and timelines. **Fig 1a** represents the ASST. Two white ceramic pots (10cm diameter, 5cm depth) were introduced at the end of the short arms. **Fig. 1b** describes the timelines used in the behavioral experiments. “A” represents the whole timeline of the experimental protocols. “B” represents the timeline of a single trial in the first experiment, where the subject waited 30 seconds in the starting area, and after removing the sliding door, following 120 seconds to explore the entire apparatus, find the rewarded pot, and retrieve the chocolate reward. “C” represents the timeline in the second experiment, where the subject waited 30 seconds in the starting area with optogenetic stimulation, and after removing the sliding door, following 30 seconds to explore the apparatus while optogenetic stimulation was still on, and further 90 seconds to find the correct pot and retrieve the chocolate reward. Finally, **fig 1c** represents the ASST apparatus with the optogenetic stimulation setup.

**Fig 2:**
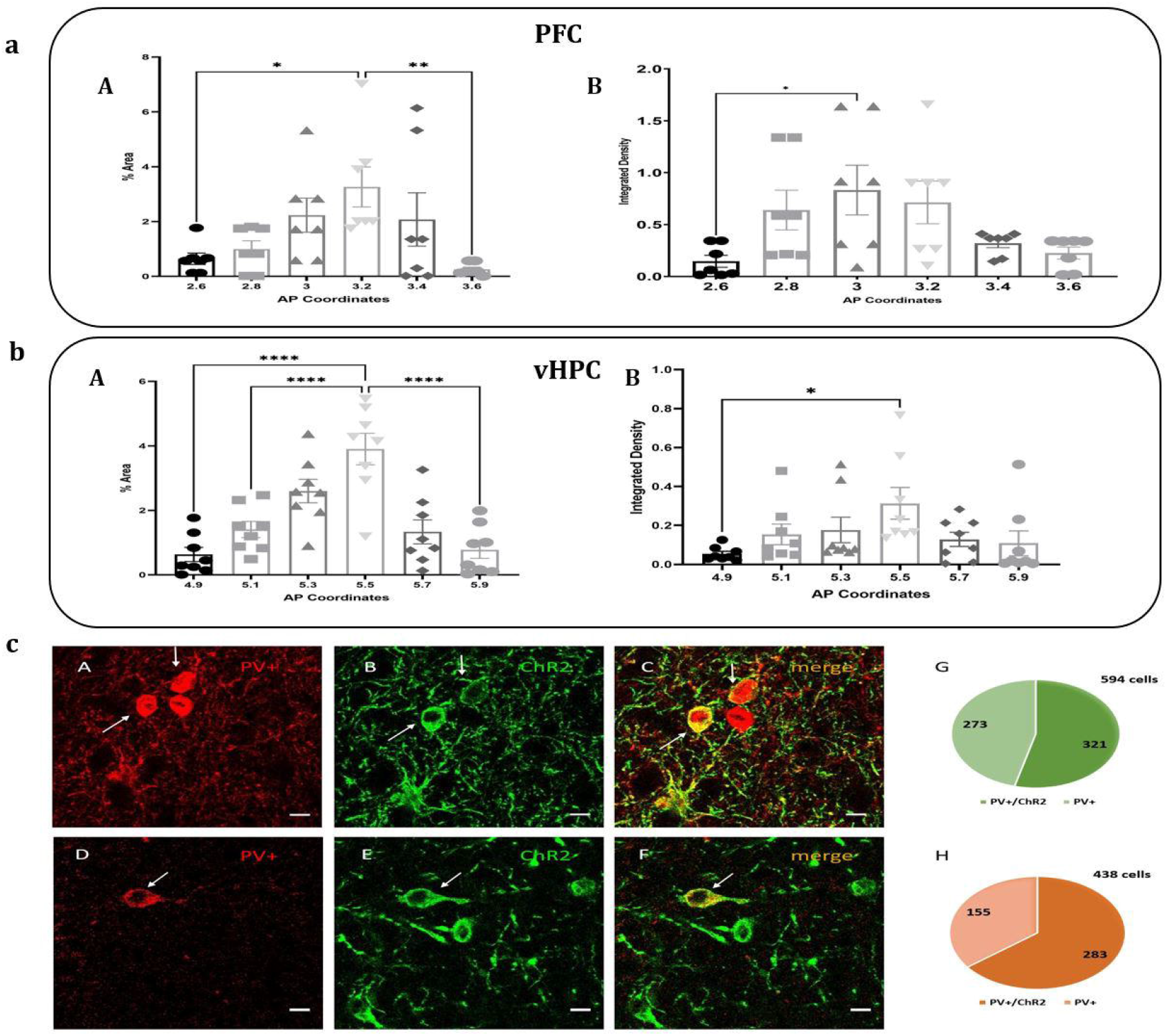
Representative example of the sectioning for PFC and vHPC regions. **Fig 2a** and **b** represent the PFC section expressing ChR2 (**a**, in green) and the vHPC section expressing ChR2 (**b**, in green). Dotted lines and the arrow represent the position of the optic fiber. The white bars in the top left corner represent the image scale (250μm in **a** and 500μm in **b**). Abbreviations: PFC: Cg1 (cingulate cortex 1); IL (infralimbic cortex); PrL (prelimbic cortex). vHPC: CA1 (Cornus hammonis 1); CA3 (Cornus hammonis 3); Py (pyramidal cells layer); VS (ventral subiculum).

**Fig 3:**
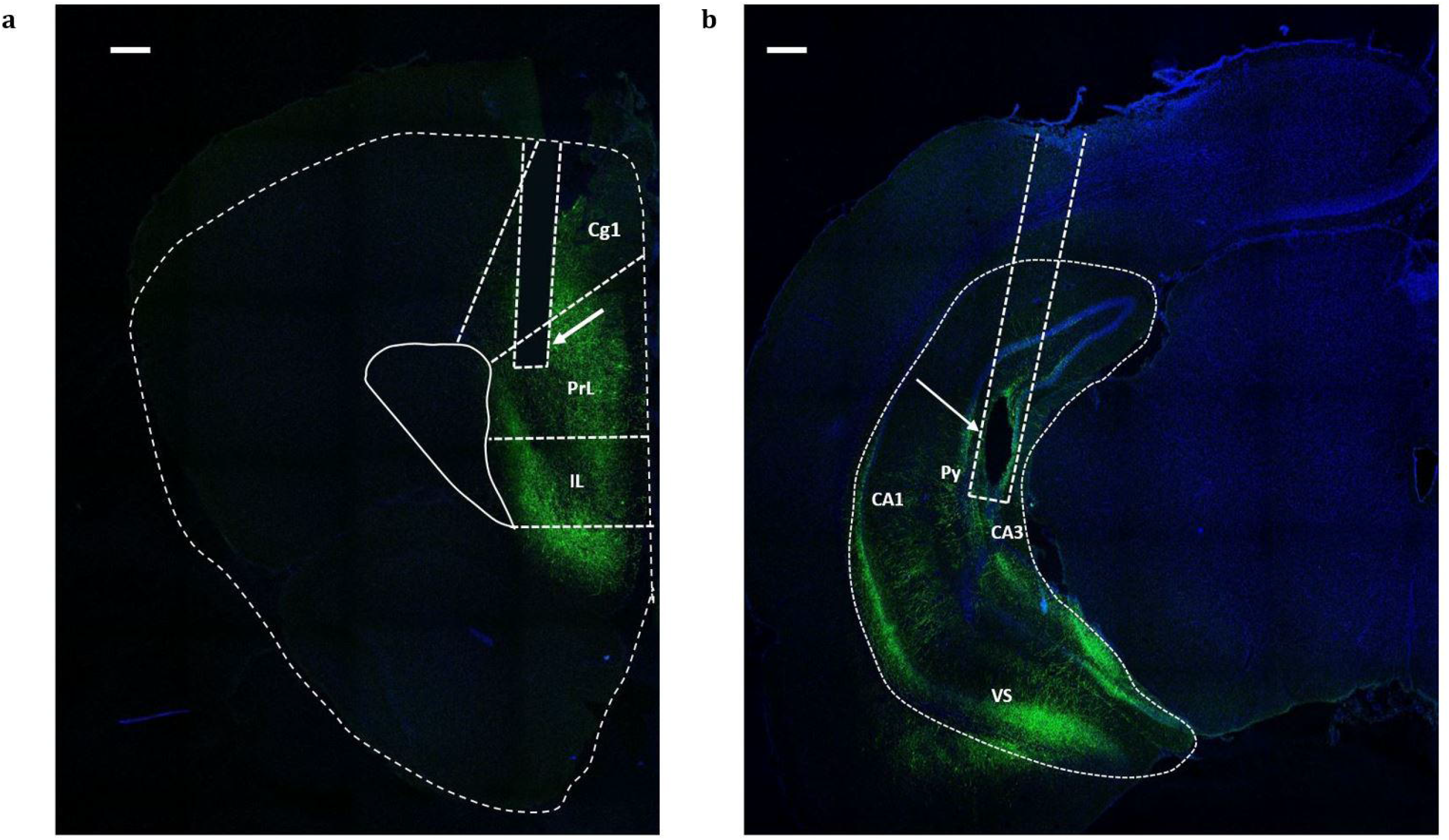
The behavioral results. **Fig 3a** shows the total number of trials before reaching the 6-CCT criterion during the CDrev, IDSrev, and EDS. In CDrev and EDS, MK801 revealed more trials than the NaCl group (* p-value < 0.05). The WT groups of PFC and vHPC showed a higher number of trials compared to their respective HET groups (* p-value < 0.05; ** p-value <0.01) and the NaCl group (* p-value < 0.05; ** p-value <0.01). **Fig 3b** shows the number of errors in the reverse/switching rule sessions. MK801 group showed significantly more errors in choosing the unrewarded pot (* p-value < 0.05; ** p-value <0.01). However, the optogenetic groups (WTs vs. HETs) did not show any difference. WT groups showed more omissions than MK801, yet not significant (**Fig 3c**). In **Fig 3d**, the latency to reach the positive pot is shown. Only the PFC_HET group shows a higher latency to reach the positive pot, compared to all the other groups, including the NaCl group (* p-value < 0.05; ** p-value <0.01; *** p-value <0.001).

We then evaluated the inability to reverse/switch to the relevant rule in the CDrev, IDSrev, and EDS for all the groups. A two-way RM ANOVA found effects on the groups (F [5, 40]=13, p<0.0001), and on the interaction (sessions x groups) (F [10, 80]=2.5, p=0.0125). Multiple comparisons revealed significantly higher errors in the MK801 group than the NaCl group, but only in the CDrev and EDS sessions. However, comparisons within the optogenetic groups did not reveal significant differences in the number of errors in any session (Fig 3b). It is worth noting that in CDrev and EDS, the HET groups showed fewer errors than the WT groups. An overall lower number of errors in the WT groups with respect to the correspondent MK801 group was due to intrinsic differences among the animal batches used for the different experiments. Moreover, further analysis of “no choices’’ trials showed a higher, although a not significant number of no pot choosing in the WT groups, with respect to the MK801 group (Fig 3c), suggesting a possible higher influence of the MK801 on the WT groups than in the MK801 group.

Concerning the latency to reach the positive pot, seconds were recorded for each correct trial and averaged in the reverse/switching rule sessions for all the groups. Therefore, a two-way RM ANOVA showed effects for the groups (F [5, 40]=21.70, p<0.0001), the sessions (F [1.907, 76.27]=9.504, p=0.0003), and the latency factor (F [40, 80]=1.770, p=0.0154), but not in the interaction (sessions x groups) (F [10, 80]=0.5571, p=0.8438). Multiple comparisons showed a significantly higher latency to reach the positive pot in the PFC_HET group with respect to PFC_WT, MK801, and NaCl groups (Fig 3d). Notably, the latency to reach the positive pot was unaffected in the vHPC_HET group compared to the vHPC_WT group. These results suggest an effect of the optogenetic stimulation of PFC PV+ interneurons on the increased driving motivation to intake a food reward possibly induced by a single MK801 injection.

### Immunohistochemistry

One-way ANOVA for the integrated density of fluorescence and % area in PFC revealed a significant role of the transduction (% area: F [5, 36] = 3,876, p= 0.0065; integrated density: F [5, 36] = 3,369, p= 0.0134). Multiple comparisons showed a significant difference between the caudal and rostral slices and the optic fiber position (% area: AP 2.6 vs 3.2, p=0.0311; AP 3.2 vs 3.6, p=0.0091; integrated density: AP 2.6 vs 3, p=0.0379) (Fig 4a). In the vHPC, a one-way ANOVA for the % area and the integrated density of fluorescence revealed a significant role of the transduction only in terms of % of transduced area (% area: F [5, 42] = 13.67, p<0,0001; integrated density: F [5, 42] = 2.407, p= 0.0524). Multiple comparisons showed a significant difference between the caudal and rostral slices and the optic fiber position (% area: AP 4.9 vs. 5.5, 5.1 vs. 5.5, and 5.5 vs. 5.9 p<0,0001; integrated density: AP 4.9 vs. 5.5 p=0.0270) (Fig 4b). These results suggest that the surgical strategy that included viral transduction and optic fiber implant was successful and trustworthy throughout the experiments.

**Fig 4:**
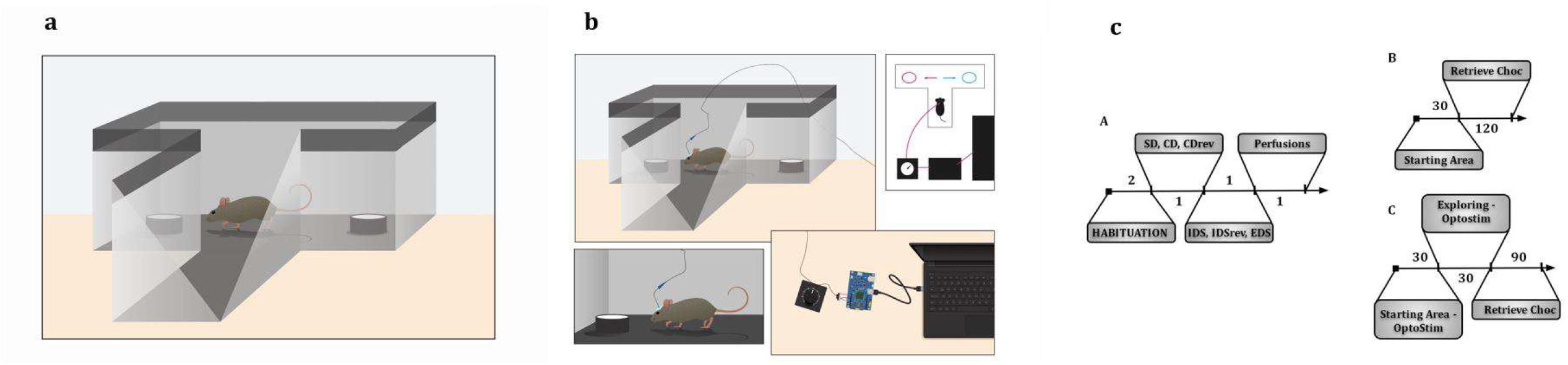
The extent of colocalization of ChR2 on PV+: **Fig 4a** shows the results of the % area (A) and the integrated density (B) of fluorescence in the six adjacent sections of the PFC, covering 1mm (+2.6/+3.6 mm AP to bregma, with the fiber implant at +3.2 mm). Significant levels of fluorescence were seen in the optic fiber position compared to the other more anterior/posterior slices (* p-value < 0.05; ** p-value <0.01). **Fig 4b** shows the results of the % area (A) and the integrated density (B) of fluorescence in the six adjacent sections of the vHPC, covering 1mm (−4.9/-5.9 mm AP to bregma, with the fiber implant at -5.5 mm). Significant levels of fluorescence were seen in the optic fiber position compared to the other more anterior/posterior slices (* p-value < 0.05; **** p-value <0.0001). **Fig. 4c** shows representative cells expressing ChR2 (in green) onto PV+ interneurons (in red) in both PFC (A, B, C) and vHPC (D, E, F) are shown, along with pie charts with total numbers of PV+ body cells detected in the six examined slices for both PFC (G) and vHPC (H). Scale bars (10μm).

PV+/ChR2 colocalization was quantified in the region-of-interest (0.1×0.1 mm) around the fiber trace. Furtherly, a one-way ANOVA ruled out a significant difference between the PFC and vHPC number of cell bodies (F [5, 39] = 8,924, p<0,0001), and specifically in the averaged number of cell bodies that were transduced or non transduced (p = 0.0187) (Fig 5a). Moreover, in Fig 5b, representative cells expressing ChR2 onto PV+ interneurons in both PFC (A, B, C) and vHPC (D, E, F) are shown, along with pie charts with total numbers of PV+ body cells detected in the six examined slices for both PFC (G) and vHPC (H).

## Discussion

### The acute MK801 model of SCZ-like cognitive inflexibility

In recent years, MK801 has mainly been validated to model the cognitive dysfunctions typical of the SCZ spectrum (19-20). Indeed, it has been demonstrated that single doses of MK801 did not affect the whole behavioral procedure but only the cognitive abilities investigated (24). Accordingly, our first set of experiments showed that a single dose of MK801 induced the inability to reverse/switch the previous odor/digging medium pairings compared to the NaCl group. Furthermore, it is essential to note that our experiment maintained rats under a mild food restriction, which imparts mild chronic stress and induces deficits in reversal learning (41). Finally, our data indicate that mild chronic stress and acute systemic MK801 caused an intermediate but evident impairment in executive functions, resembling those in the SCZ spectrum.

Perseverative errors are widely considered deficits of executive functions in SCZ, and they can be used as a qualitative characterization of cognitive performance (42). Overall, we observed that the MK801 group committed more errors than the NaCl group, suggesting that the acute MK801 injections may have induced the inability to reverse/switch previous sets of associations, thus modeling cognitive inflexibility.

About the latencies to reach the correct pot, we noticed that the MK801 group had similar results as the NaCl group. Furthermore, recent findings demonstrated the role of MK801 in increasing impulsivity and the drive towards rewards (43). Therefore, we argued that MK801 acute injections might affect the impulsivity for chocolate drop assumptions.

### The role of PFC-vHPC PV+ optogenetic stimulation in the ASST

We compared the first experiments’ reverse/switching rule sessions (NaCl vs. MK801) with the optogenetic experiments. Our results showed a main effect of the optogenetic stimulation of the PV+ interneurons in both the PFC and vHPC. The results resembled those obtained by the first experiment with NaCl and MK801 groups. Furthermore, the HET groups of both the PFC and vHPC had a lower number of trials to reach the 6-CCT criterion, similar to the NaCl group (Fig 3a), showing that frequency-specific light stimulation of the PV+ interneurons in both the PFC and vHPC can counteract the compromised cognitive flexibility. The PFC and vHPC share common neurophysiological pathways playing a pivotal role in cognitive abilities, and disruption of these connections results in neuropsychiatric disorders (44). Information in large prefrontal neuronal assemblies can be synchronously transferred to the hippocampal associative networks with gamma oscillations and hippocampal theta waves (32). Moreover, GABAergic interneurons can regulate firing rate, spike timing, and synchronization, allowing for information encoding (33-35). Therefore, NMDAR hypofunction induced by MK801 in cortical and hippocampal PV+ GABA interneurons may disrupt spike synchronization. Our optogenetic experiments stimulated the PFC PV+ interneurons using a gamma-like frequency (50 Hz) and the PV+ in vHPC using a theta-like frequency (10 Hz), and we’ve been able to recover the theta/gamma synchronization previously disturbed by MK801. Our findings align with previous studies, hypothesizing that optogenetic stimulation at an appropriate frequency can counteract the PFC-gamma/vHPC-theta desynchronization induced by systemic injection of MK801. However, to our knowledge, this is the first time that a PV+ inducing re-synchronizing activity has been tested in rats on a high-level cognitive task, such as the ASST.

The number of errors in PFC and vHPC experiments generally decreased compared to the NaCl and the MK801 groups. However, in PFC and vHPC experiments, we did not observe significant differences between the HET and the WT groups. It has been suggested that Although mPFC may be involved in the initial suppression of established response sets after rule relevancy changes (45), a recent study found that optogenetic silencing of PFC did not affect the perseverative errors (46). Nevertheless, it is possible to consider general perseverative errors as a measure of perseverative behavior (47), thus defining cognitive inflexibility. This evidence suggests that our optogenetic stimulation of PV+ increased the overall ability to choose a pot.

Latency changes in the ASST can be understood as differences in speed, accuracy trade-off, or simply alterations in locomotor activity. Our experiments recorded latencies to reach the positive pots in the CDrev, IDSrev, and EDS sessions. The results showed a significantly higher latency in the PFC_HET group with respect to PFC_WT, MK801, and NaCl groups. However, the latency to reach the positive pot was unaffected in the vHPC_HET group compared to the vHPC_WT group (Fig 3d). These results suggest an effect of the optogenetic stimulation of PFC PV+ interneurons on the increased driving motivation to food reward intake and impulsivity, possibly induced by MK801 single injections (44).

### Validation of ChR2 expression and fiber position in PFC and vHPC brain areas

To validate the expression of ChR2 onto PV+ interneurons in PFC and vHPC areas and to verify the position of the optic fiber in targeted regions, we used a complex strategy based on immunohistochemical dual-labeling, the quantification of PV+/ChR2 colocalization, and the quantification of cells that were transduced/reached by light stimulation. Results showed that in both the PFC and vHPC, the % of the transduced area and the integrated density of the transduced cells were significantly higher in the target point (AP +3.2 for PFC, and AP -5.5 for vHPC), suggesting that the mechanical local injection of AAV-containing-ChR2 was successfully performed and that the optic fiber was positioned where it was likely to interact with the transduced cells.

Concerning the quantification of the PV+/ChR2 colocalization, the superposition of fluorescence images is the most prevalent method for evaluating colocalization. All biological-image analysis software implemented tools for displaying multiple-channel fluorescence images as merged color images (48). The intensity of a pixel in one channel is evaluated against the corresponding pixel in the second channel, generally producing a correlation coefficient. Unfortunately, Pearson’s correlation coefficient is not sensitive to mean signal intensities or range differences. However, another coefficient using a proportion of the colocalizing pixels fluorescence appears to be reliable (Manders split coefficient, 49). Therefore, a statistical test is performed by randomly scrambling hundreds of times the blocks of pixels/voxels in one image and then measuring the correlation of this image with the other image (50). This test returns a high level of correlation measurement, surpassing the manual images thresholding methods, which is subjective and therefore poorly reliable. Our colocalization analysis showed a high rate of colocalization among PV+ interneurons with ChR2, suggesting that the PV+ were successfully activated by ChR2 when reached by light. Therefore, this result validates previous behavioral data demonstrating that appropriate optogenetic stimulation of PV+ interneurons in either the PFC or vHPC is key to re-establishing cognitive flexibility after acute MK801 injections.

In conclusion, in this study, we determined a new animal model of SCZ-like cognitive inflexibility, using acute systemic injections of MK801 in the reverse/switching rule sessions; and we evaluated the ability of frequency-specific optogenetic stimulations of PV+ interneurons in two critical brain structures (the PFC and vHPC) to counteract the MK801-induced cognitive inflexibility. Furthermore, in line with previous studies, this work outlined the importance of hippocampal-prefrontal communication as a critical link in a cognitive control mediating network. Finally, this study will serve as a starting point for further investigations to expand knowledge on what specific processes the vHPC and PFC perform in this network.

## Supporting information

Suppl Mat

## Acknowledgments

The funding for designing and writing the manuscript was provided to EP by ESF project CZ.02.2.69/0.0/0.0/19_074/0016409. KH was supported by the Charles University project GA UK 377521. AS and JS were supported by Czech Science Foundation (GACR) grants 19-03016S, 20-00939S, and 21-16667 K, respectively. Institutional support for IPHYS was provided by RVO: 67985823. We thank Alessandro Ribaldo for Fig. 1a and c. We also thank Dr. David W. Hardekopf for his proofreading services.

