## Supplementary material for "The role of optogenetic stimulations of parvalbumin-positive interneurons in the prefrontal cortex and the ventral hippocampus on an acute MK801 model of schizophrenia-like cognitive inflexibility": Suppl Mat

### Supplementary Materials

#### *The attentional set-shifting task*

The apparatus consisted of a new T-maze version (37) (long arm, 29x15x25cm; short arms, 29x15x25cm). A sliding door separated the long arm as a starting area from the short arms. Two white ceramic pots (10cm diameter, 5cm depth) were introduced at the end of the short arms (Fig 1a). The pots were defined by cues along two dimensions: odor (relevant) and digging medium (not relevant). A bait in the form of a chocolate drop (∽150 mg, Juweel, Germany) was placed at the bottom of the rewarded (positive) pot and buried in the medium. The positive pot was then paired with a relevant odor (e.g., oregano) and a not relevant digging medium (e.g., straw). The not rewarded (negative) pot was paired with another not relevant odor (e.g., parsley) and another not relevant digging medium (e.g., shredded paper). Table 1 describes the relevant/not relevant dimensions that were rewarded/not rewarded according to the task session (Tab 1).

Before starting the ASST, chocolate drops (5 g) were placed in the home cages to avoid novelty effects. During habituation (two days), rats were trained to dig in the pots, obtaining rewards reliably. Subsequently, underwent a simple discrimination task (SD) where a simple association (odor/chocolate) was taught, reaching a criterion of six consecutive correct trials (6-CCT). In detail, the rat was gently put in the starting area for 30 secs. Then, the sliding door was removed, and the rat had 2 mins to find the positive pot and dig up and retrieve the chocolate drop. After SD, rats were exposed to increasingly challenging tasks, including compound discrimination (CD) and CD reversal (CDrev). The CD had the two stimulus dimensions, odor and digging medium. Therefore, rats learned associations between the relevant odor, not relevant digging medium, and the food reward. Once the 6-CCT was reached in the CD session, MK801 and NaCl were injected i.p., respectively, and the rats were moved to the cages. After 30 mins, the rats performed CDrev. In the CDrev, the positive pot association was reversed based on the relevant dimension. On the fourth day, rats were exposed to an intradimensional shift (IDS), an IDS reversal (IDSrev), and an extradimensional shift (EDS). The IDS was similar to CD, except for different odors and digging mediums. Again, following IDS, rats of both groups were injected with either MK801 or NaCl i.p., respectively, and then tested for IDSrev and EDS. In the IDSrev, the positive pot association was reversed based on the previous IDS. In the EDS, the relevant dimension was switched from odor to digging medium, assuming a newly relevant digging medium/chocolate combination. Fig 1b shows the protocol timeline used in the experiments (Fig 1b). We tested relevancy shifting from odor to digging medium to better compare reversal and switching rule sessions (38). In this set of experiments, perseverative errors were scored when a rat continued to use the previously relevant but currently irrelevant rule. No choice was scored when the rat did not choose any pot after 2 mins. Inter-trial shifts were used in each session to ensure that the subjects had chosen the positive pot based on the relevancy rule. The positive pot was pseudo-randomly switched between the two short arms to avoid side preference effects during the whole protocol. The ASST testing typically took 75-90 min per rat per day to complete.

#### *Surgeries and viral injections*

Rats were anesthetized with isoflurane (5% induction, 2% maintenance) and locally by subcutaneous injections of mesocaine (0.2 ml). For viral injections, the following flat skull coordinates relative to the bregma were used, according to the rat atlas (39): PFC (AP +3.2, ML ±0.6, DV -4/-3), vHPC in 10° angle (AP -5.5, ML ±5, DV -5.8/-5.4). ChR2 was infused (0.500 µl each spot at 0.050 µl/min flow rate) using a Hamilton syringe (5 µL, #87943, Hamilton, USA) attached to an infusion pump (UMP3 with SYS4 Micro-controller, World Precision Instruments). The needle was left in place for 5 min to allow for diffusion. After viral injections, a cannulated optic fiber (400 mm diameter, 0.39 NA, Thorlabs, Germany) was placed at the coordinates described above, with DV -3.5 mm for the PFC and DV -5.6 mm for the vHPC, from the brain surface. Two anchoring screws were placed near the optic implant. A mix of cyanoacrylate (Loctite SuperBond) and methacrylate glue (dental cement, Duracrol) was applied to ensure sufficient fixation of the optic implant throughout the whole experiment. Postoperative care was provided by adding antibiotics and analgesics (neomycin and ibuprofen) to the drinking water. The rats were checked daily and left to recover for three weeks. The behavioral tasks were not started until the animals showed no discomfort when manipulating the implants. Furthermore, the three-week recovery allowed ChR2 to express the PV+ interneurons in the targeted areas.

#### *Behavioral assessment*

Rats had *ad libitum* access to food and water. However, one week before and during the behavioral protocols, the animals were food-restricted and maintained at 85–90% of their body weight.

Fig 1c represents the behavioral setting with the optogenetic setup (Fig 1c). In the CDrev, IDSrev, and EDS, the rats were attached to a patch cable connected to a fiber-coupled LED (M470F3, 470 nm, Thorlabs, Germany) and an LED Driver (LEDD1B, Thorlabs, Germany) and gently put in the starting area. After the first 30 secs, the sliding door was removed, and the rats had 2 mins to choose and dig to retrieve the chocolate. The light was turned immediately off when the rat retrieved the reward from the positive pot and was gently confined into the starting area. Otherwise, after 30 secs, the light was automatically shut off, and the rat was left to choose a pot for another 90 secs. The time to reach and retrieve the reward was measured, along with incorrect or no choices. Subsequent trials followed the same procedure until the animal reached the 6-CCT. A day after ending the ASST, perfused brains were fixed in 4% paraformaldehyde for 24 hours, soaked in 30% sucrose solution, and stored at -80°C for immunohistochemical assays.

#### *Immunohistochemistry*

Coronal sections (50 μm thickness) of PFC and vHPC areas were cut using a cryostat (Leica CM1950) to verify viral expression in the optic fibers and the correct position in the PFC and vHPC. Brain sections of both HET and WT groups were collected and cryopreserved. In addition, six adjacent sections were collected (every 200 μm, ca), covering 1 mm anteroposterior distance to visualize the optic fiber position and the viral expression in both PFC and vHPC transduced areas. Fig. 2a and b show a representative example of the sectioning for PFC and vHPC regions, with the ChR2 expression around the fiber trace (Fig 2a, b).

The collected sections were washed in TBS, permeabilized with TBS and Triton-X 100 1.2% (20mins, RT), and blocked with TBS and normal bovine serum (NBS) 5% (60 mins, RT). Sections were then incubated in rabbit polyclonal antibodies against PV (Abcam, ab11427,1:500) and against chicken anti-green fluorescent protein (GFP) antibody (Abcam ab13970, 1:500) in a TBS solution containing 2% NBS and 0.2% Triton X-100 (4°C, overnight). The sections were then washed and incubated in donkey anti-rabbit Alexa Fluor 594 (Invitrogen A32754, 1:500) and anti-chicken Alexa Fluor 488 (Invitrogen A11039, 1:500) secondary antibodies (2 hrs, RT). Brain sections were then washed in TBS, mounted onto gelatin-coated slides, and coverslipped with Vectashield antifade mounting medium containing DAPI (Vector Laboratories, #H1200). Finally, images from the brain sections were acquired with a confocal microscope (Leica SP8X) and analyzed using Fiji and ImageJ software.

#### *Data Analysis*

The ASST is a standardized task that measures the rat's ability to reverse/switch the relevancy of a discrimination rule to solve the task and be rewarded. Here, we measured the total number of trials to reach the 6-CCT criterion in each session. Moreover, since perseverative errors are considered a measure of inability to disengage from an old rule, the incorrect choices performed by the animals were scored during the reverse/switching rule sessions. Furthermore, latencies to reach the positive pots were recorded during the reverse/switching rule sessions since latency changes can be understood as differences in speed, accuracy trade-off, or simply alterations in locomotor activity (40). Therefore, all these parameters were measured in all experiments.

*Viral expression and optic fiber position:* the quality, the extent, and the functionality of the expression of ChR2 onto PV+ interneurons were evaluated in PFC and vHPC. Acquisition of confocal images at 10x was made from each of the six adjacent sections from each of the HET brains to locate the fiber traces and evaluate the density of the ChR2 expression around the region of the fiber traces, using the integrated density and the % area of fluorescence parameters. Secondly, we quantitatively measured the colocalization of ChR2 onto PV+ interneurons. Therefore, confocal images at 40x have been acquired for every six slices for each subject from PFC and vHPC experiments. A region-of-interest (0.1x0.1mm) around the fiber trace has been used to analyze each slice where the fiber trace was present. Manual counting of PV+ cell bodies (in red) in the region-of-interest and PV+/ChR2 colocalization quantification have been performed. Table 2 shows the results of the manual counting of PV+ body cells and the percentage of transduced vs. non-transduced cells in PFC and vHPC (Tab 2).

The intensity and the density of the PV+/ChR2 colocalizations were quantified using plugins in Fiji and ImageJ (Coloc2, JacoP) (Tab 3). The intensity and density of the colocalizations are reported as correlation coefficients (Pearson's coefficient, Manders's coefficient, and Costes's coefficient) from the superimposition of fluorescence images, using pixel matching colocalization analyses. The Coloc2 analysis reported high levels of colocalization intensity in the cell bodies in the region-of-interest in both the PFC and vHPC. The JacoP analysis reported high levels of colocalization density in the region-of-interest in both the PFC and vHPC.

**Table 1: Order of discriminations.**

| **Session** | **Relevant Dim.** | **Not Relevant Dim.** | **Rewarded** | **Not Rewarded** |
|---|---|---|---|---|
| **SD** | **ODOR** | **NONE** | Oregano | Parsley |
| **CD** | **ODOR** | **MEDIUM** | Oregano/Grinded<br>-----------------------------<br>Paper Oregano/Straw | Parsley/Straw<br>------------------------------<br>Parsley/Grinded Paper |
| **CDrev** | **ODOR** | **MEDIUM** | Parsley/Straw<br>------------------------------<br>Parsley/Grinded Paper | Oregano/Grinded<br>Paper ---------------------<br>--------- Oregano/Straw |
| **IDS** | **ODOR** | **MEDIUM** | Marjoram/Floral Foam<br>------------------------------<br>Marjoram/Sponge | Basil/Sponge<br>------------------------------<br>-<br>Basil/Floral Foam |
| **IDSrev** | **ODOR** | **MEDIUM** | Dill/Stones<br>------------------------------<br>Dill/Clay | Rosemary/Clay<br>------------------------------<br>-<br>Rosemary/Stones |
| **EDS** | **MEDIUM** | **ODOUR** | Sand Paper/Fennel<br>------------------------------<br>Sand Paper/Thyme | Cork/Thyme<br>------------------------------<br>-<br>Cork/Fennel |

Examples of combinations into stimulus pairs are shown for a rat shifting from odor to digging medium. Inter-trial shifts were used in each session to ensure that the subjects had chosen the positive pot based on the relevancy rule. The combination of exemplars into rewarded /not rewarded stimuli and their left-right position of presentation in the T-maze was a pseudorandom series determined in advance.

**Table 2: Total number of PV+ body cells.**

| Total PFC | Transduced | % | Not Transduced | % |
|---|---|---|---|---|
| 594 | 321 | 54% | 273 | 46% |
| Total vHPC | Transduced | | Not Transduced | |
| 438 | 283 | 64.6 % | 155 | 35.4% |

**Table 3: The intensity and density of the colocalization reported as correlation coefficients.**

| PFC | | | |
|---|---|---|---|
| Coloc2 (Intensity) | Costes | Manders | Pearson |
| | 0.938 | 0.803 | 0.70 |
| JACOP (Density) | Costes | Manders | Pearson |
| | 0.830 | 0.670 | 0.350 |
| vHPC | | | |
| Coloc2 (Intensity) | Costes | Manders | Pearson |
| | 0.926 | 0.645 | 0.71 |
| JACOP (Density) | Costes | Manders | Pearson |
| | 0.890 | 0.850 | 0.350 |
